# Transport kinetics across interfaces between coexisting liquid phases

**DOI:** 10.1101/2024.12.20.629798

**Authors:** Lars Hubatsch, Stefano Bo, Tyler S. Harmon, Anthony A. Hyman, Christoph A. Weber, Frank Jülicher

## Abstract

Biomolecular condensates provide compartments that organize biological processes in the cell. Spatial organization of condensates relies on transport across phase boundaries. We investigate the kinetics of molecule transport across interfaces in phase-separated mixtures. Using non-equilibrium thermodynamics we derive the interfacial kinetics, describing the movement of the interface, and of molecular transport across the interface. In the limit of a thin interface, we show that breaking local equilibrium at the interface introduces a local transport resistance. Subsequently, using a continuum approach, we explore two physical scenarios where such a resistance emerges: a mobility minimum and a potential barrier at the interface. These scenarios lead to the same effective transport model in the sharp interface limit, characterized by a single interface resistance parameter. We discuss experimental conditions for which interface resistance could be observed and how these scenarios could be distinguished.

## I. INTRODUCTION

In biological cells, membraneless condensates can be understood as droplets of coexisting liquid phases [1]. These condensates form biochemical compartments in the cell. Such condensates can dynamically form and dissolve, and they need to exchange molecules with their surroundings to enable biochemical processes [2]. The transport kinetics across condensate interfaces can thus impact cellular function and dysfunction. To understand how biomolecular condensates affect the dynamics of key biological processes, it is important to quantify these kinetics in the coexisting phases and across phase boundaries [2].

Experimental assays such as fluorescence recovery after photobleaching (FRAP) and single-molecule localization microscopy (SMLM) can be used to study molecular transport inside the phases and across interfaces of biological condensates. This transport is characterized by kinetic parameters such as diffusion coefficients and thermodynamic properties such as partition coefficients [3–10].

Phase boundaries have been investigated in different systems with different methods, from *in vivo* condensates in the cell [11], to aqueous two-phase systems in chemistry and engineering [12]. For example, in the cell nucleus, protein diffusion was quantified close to the heterochromatin-euchromatin border indicating a slower protein diffusion at the phase boundary compared to the surrounding phases [11]. Other *in vivo* work quantified single molecule trajectories moving across the phase boundary of viral replication compartments in the nucleus [13] and the confinement effects that biomolecular condensates have on their constituents [14].

In reconstituted protein condensates, Taylor et al. found a low diffusion coefficient inside condensates when fitting a transport model based only on two diffusion coefficients and the partition coefficient [5]. They suggested that this apparent slow diffusion might be an artifact of the simple transport model, and thus incorporated an interfacial resistance parameter. Such an effect had been introduced in Ref. [15, 16] in the context of ripening kinetics [17]. However, the magnitude of this effect remains unclear due to the challenge of accurately quantifying partition coefficients by fluorescence [18], which in turn makes it difficult to create reliable fits to theoretical models. Subsequent work on other *in vitro* protein condensates found good agreement of a theory without interface resistance with experimental data [3].

In aqueous two phase systems, made, for example, of polyethylene glycol and dextran, phase boundaries can be observed using microfluidics setups. Applying an electric field across the phase boundary, one can observe the motion of several model proteins or DNA across the phase boundary [7, 8]. Under certain conditions, this resulted in a transient accumulation of molecules at the liquid-liquid interface, slowing interfacial transport. This slowdown was attributed to the interplay between the applied electric field and the Donnan potential across the interface [12].

Recent theoretical work has discussed possible physical origins of interface resistance. For example, interfacial charge-separation was suggested to influence interfacial transport [19]. There is also numerical evidence for interface resistance from studying a coarse-grained model of stickers and spacers [6]. Taken together, the extent to which additional effects at the phase boundaries have to be considered requires both theoretical and experimental clarification.

Here, we present a general theoretical framework, based on irreversible thermodynamics, to capture interfacial transport kinetics in multi-component phase-separated systems and discuss its application to experiments. We start by developing our approach for a binary mixture in a sharp interface limit, taking into account dissipation within the interface. Dissipation arises from differences of chemical potentials across the interface, corresponding to a phase boundary that is not at phase equilibrium. This situation is associated with interface resistance, slowing the movement of phase boundaries and molecular transport across phase boundaries. Our framework naturally extends to multiple molecular species, and we show how interface resistance influences the exchange kinetics of labeled and unlabeled molecules across a phase boundary. We then generalise our approach to a continuum theory where the interface has a finite width. We demonstrate that a potential barrier or a reduction in diffusivity within the phase boundary can contribute to interface resistance. Finally, we propose experimental settings *in vitro* and comment on the necessary optical resolution to detect signatures of interfacial kinetics associated with resistance and distinguish the underlying physical mechanisms.

## II. TRANSPORT ACROSS AN INTERFACE IN A SHARP-INTERFACE LIMIT

### A. Interface dynamics in a binary mixture

This section aims to derive the fluxes of condensate and solvent (subscript “s”) molecules at the phase boundary between a dense phase (subscript “in”) and a dilute phase (subscript “out”). To this end, we consider a sharp interface model where the interface can move.

We start from a binary mixture with free energy density

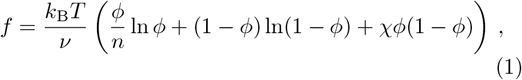

where *ϕ* denotes the volume fraction of the condensate molecule. We consider an incompressible system with a constant molecular volume of the solvent *ν*, and a constant *n* denoting the molecular volume fraction between the condensate molecule and solvent. Moreover, *χ* is an interaction parameter characterizing the interaction between both components. The solvent volume fraction is given by *ϕ*_*s*_= 1 − *ϕ*. The exchange chemical potential 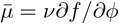 thus reads

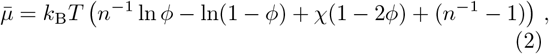

and the osmotic pressure 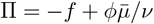 is given by

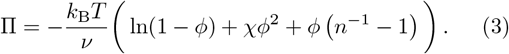

The chemical potential of the condensate molecule is 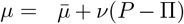, while the solvent chemical potential reads *μ*_*s*_ = *ν*(*P* − Π), where *P* denotes the pressure.

At phase equilibrium between the phases “in” and “out”, we have 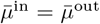 and for the Laplace pressure, we obtain ΔΠ = Π^in^ − Π^out^ = Δ*P*, where Δ*P* = *γ/R*, is the Laplace pressure and *γ* denotes the surface tension.

The dynamic equation for condensate molecules and the solvent are:

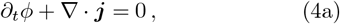

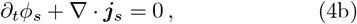

Incompressibility implies that fluxes of solvent and condensate molecules are equal and opposite in each phase: ***j***_**s**_ = − ***j***.

Since the volume fraction *ϕ* jumps discontinuously at the interface when the interface moves, the flux ***j*** changes discontinuously, too. In the reference frame moving with the interface, the magnitude of the normal flux *j* = ***n*** · ***j*** is continuous across the interface, where ***n*** is the outward normal vector on the interface. Therefore, we have

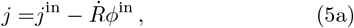

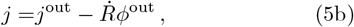

where 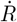 is the outward normal interface velocity, *j*^in^ = ***n*** · ***j*** ^in^ and *j*^out^ = ***n*** · ***j***|^out^ are the normal fluxes evaluated inside and outside at the interface in the lab frame. The interface velocity in the lab frame (Fig. 1A) obeys

**Figure 1.**
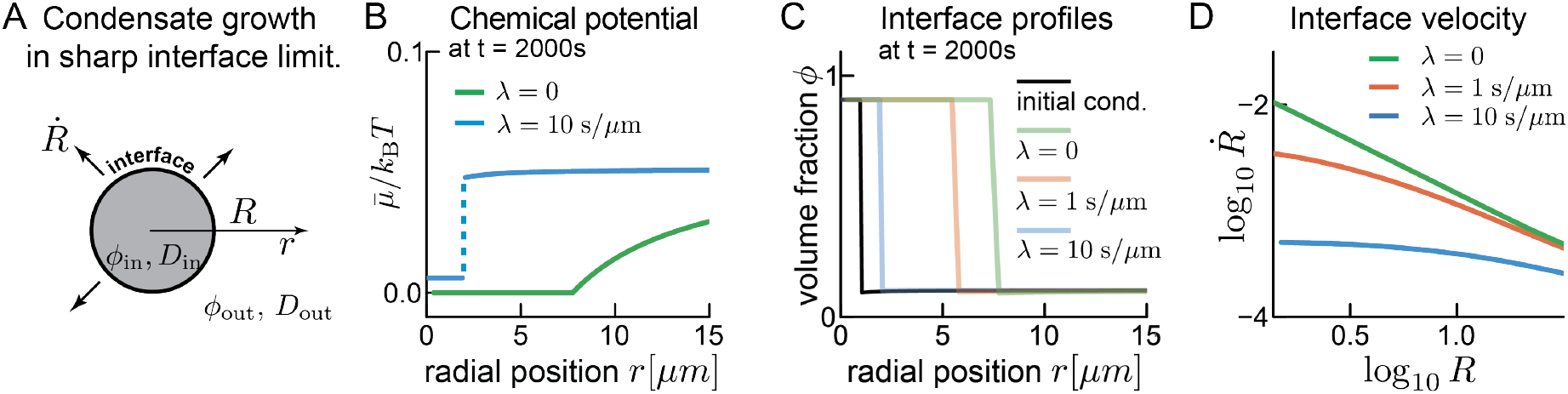
Influence of interface resistance on droplet growth. **(A)** The sharp interface limit considers two spatial domains (volume fractions *ϕ*^in^ and *ϕ*^out^, with diffusion coefficients *D*^in^ and *D*^out^) separated by a boundary, where the volume fraction jumps. In the spherically symmetric case considered here, the phase boundary moves until equilibrium is reached. **(B)** The chemical potential at the interface can be continuous (green) or discontinuous (blue, jump indicated by dashed line), resulting in different boundary kinetics (see panels (C) and (D)). **(C)** Snapshots of the volume fraction of condensate material at *t* = 2000s for different interface resistance. The droplet grows due to a diffusive flux from the right side. Systems size, 100 *μm*. We use *λ* = *λ*_s_. For numerics see appendix B**(D)**: Interface resistance plays a larger role in smaller droplets. In larger droplets diffusive flux dominates.

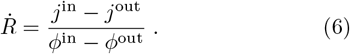

Analogously to Eq. (5a), we can write the flux of solvent in the frame of the interface, 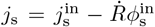 and 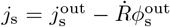. Together with Eq. (5a), we find that the interface velocity in the lab frame is opposite to the sum of fluxes of solvent and droplet material in the interface frame:

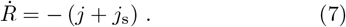

Since concentrations at the interface, in general, change discontinuously, we propose the chemical potentials *μ*^in^ and *μ*^out^ could also differ at the interface (Fig. 1B). In this case, since the interface is considered as sharp, differences between the two chemical potentials occur at the same position, suggesting a similarity to chemical potential differences between chemical states. Irreversible thermodynamics suggests a non-linear relation between chemical potentials and the fluxes *j* and *j*_*s*_ at the interface [20]:

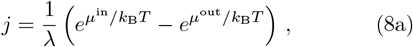

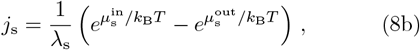

which reads upon linearization:

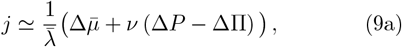

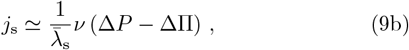

where we introduce the difference of chemical potential 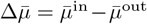, the difference in osmotic pressure ΔΠ = Π^in^ − Π^out^ and the pressure difference Δ*P* = *P* ^in^ − *P* ^out^ at the interface. Moreover, *λ* and *λ*_*s*_ are kinetic coefficients characterizing the resistance of the interface to approach phase equilibrium with respect to condensate molecule and solvent, respectively. For *λ* = 0 and *λ*_s_ = 0, we recover *μ*^in^ = *μ*^out^ and 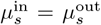, which is the case of local phase equilibrium at the interface.

If we set *λ* = *λ*_*s*_ and vary their magnitude, we find that larger interface resistance results in slower droplet growth (Fig. 1C). The interface velocity 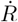is dominated by interface resistance at small radii, however, as the droplet grows, its interface velocity approaches the limit in which diffusive flux dominates interface resistance (Fig. 1D).

### B. Dynamics of labeled and unlabeled molecules

Experimentally, the dynamics of droplets are often studied using fluorescence microscopy. A commonly used imaging technique is fluorescence recovery after photo-bleaching (FRAP). In such experiments, molecules in a defined region are bleached, and the exchange kinetics with the surroundings are studied. Thus, FRAP enables visualization of molecular motion even in steadystate systems. We thus distinguish between labeled and unlabeled condensate molecules, or equivalently, unbleached and bleached molecules in a FRAP experiment (see Fig 2A). In our theory, the key assumption is that labeled and unlabeled molecules show the same physical behaviour (same molecular volume *nν* and interaction parameter *χ*). However, being able to distinguish both gives rise to a mixing entropy [3, 4, 21].

**Figure 2.**
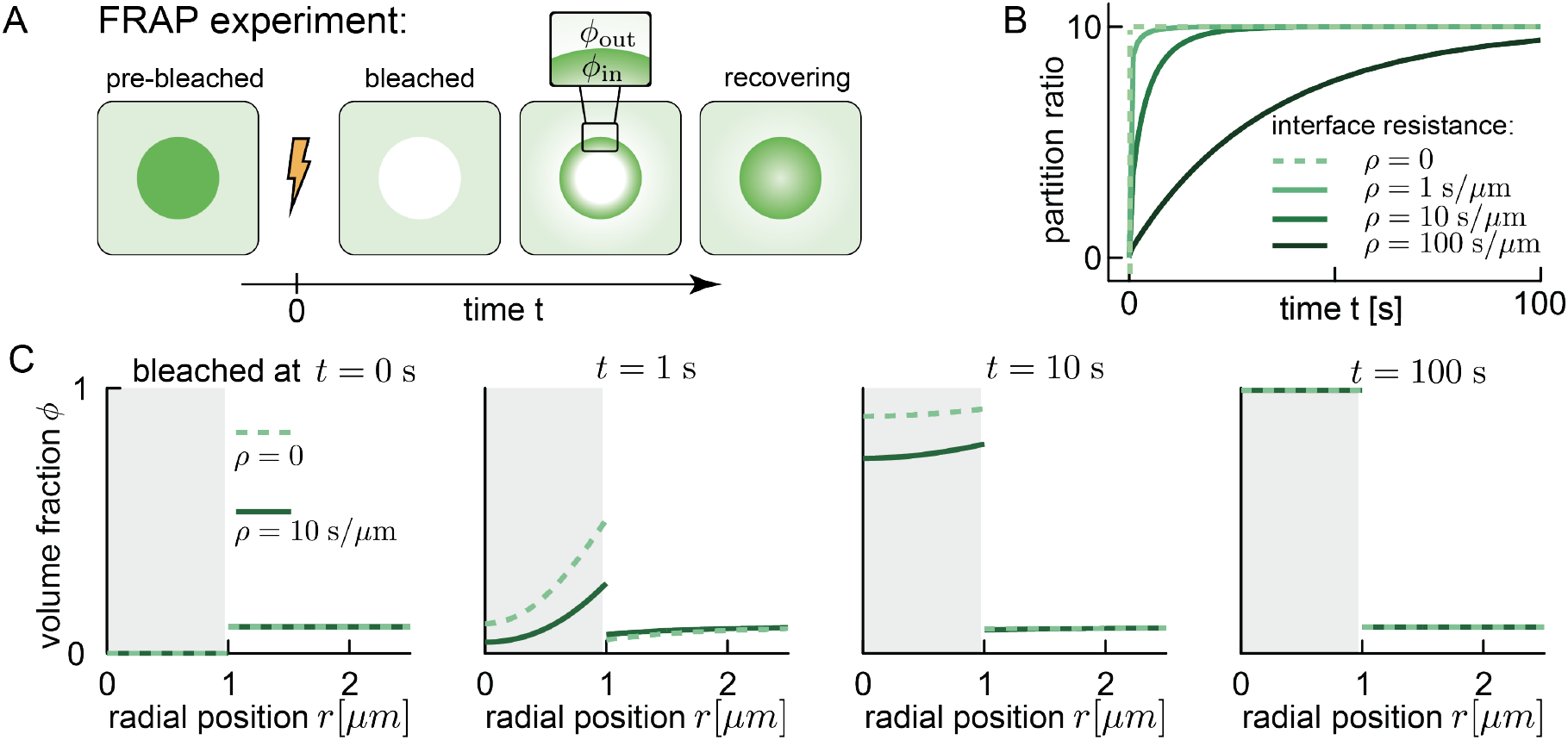
Influence of interface resistance on FRAP recovery. **(A)**: In a typical FRAP experiment, the droplet is close to equilibrium prior to the experiment. The fluorescent dye labeling the condensate material gets bleached, and is therefore not visible in fluorescence microscopy (unlabeled, white). Subsequently, the exchange of labeled and unlabeled material across the phase boundary can be observed. **(B)**: The ratio between the concentrations right inside and outside (see panel (A), *ϕ*_in_, *ϕ*_out_) starts at zero (postbleach). Without interface resistance (grey dashed line), the ratio jumps immediately to the equilibrium partition coefficient Γ^*∗*^. For the case of interface resistance, the concentration ratio approaches the equilibrium value continuously. **(C)** Time course of FRAP recovery with and without interface resistance. Initial conditions and equilibrium values are identical, but with resistance recovery is slowed down.

We denote the volume fraction of labeled and unlabeled molecules by *ϕ*_1_ and *ϕ*_2_. The total volume fraction of condensate molecules is given by *ϕ* = *ϕ*_1_ + *ϕ*_2_. The free energy then reads [3]:

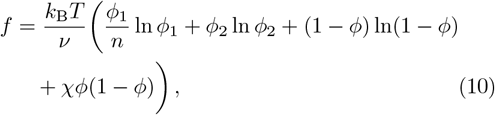

and exchange chemical potentials 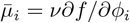 are

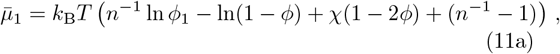

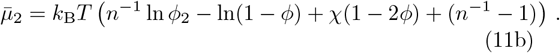

The osmotic pressure Π = −*f* + *ϕ*_1_*μ*_1_*/ν* + *ϕ*_2_*μ*_2_*/ν* reads

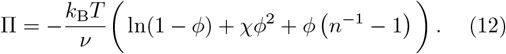

where we used 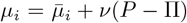. We note that while the exchange chemical potentials differ when distinguishing labeled and unlabeled molecules to the binary system (Eq. (2)), the osmotic pressure remains unchanged (Eq. (3)).

We can now follow a similar strategy as for the binary mixture and obtain the fluxes at the interface. The dynamic equation for labeled and unlabeled condensate molecules reads

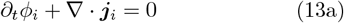

with *i* = 1, 2. For the solvent, we have

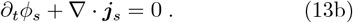

Incompressibility implies that fluxes of solvent and the labeled and unlabeled condensate molecules balance: ***j***_s_ = −***j***_1_ −***j***_2_. The sum of the labeled and unlabeled fluxes gives the flux of total condensate molecules and solvent molecules as discussed in the last section: 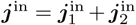 and 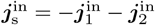.

In the reference frame moving with the interface, the magnitude of the normal fluxes *j*_*i*_= ***n*** · ***j***_*i*_ (*i* = 1, 2) and *j*_s_ = ***n*** · ***j***_s_ are continuous across the interface, where ***n*** is the normal vector to the interface. In analogy with Eq. (5a), we have

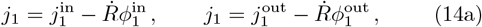

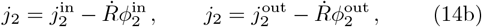

where *j*_1_ and *j*_2_ are the fluxes in the reference co-moving with the interface of labeled and unlabeled molecules, respectively. Note that 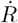 is the interface velocity of the condensate composed of labeled and unlabeled condensate molecules. Moreover, 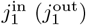 is the normal flux of labeled molecules at the interface, evaluated just inside (outside) the condensate. The solute flux obeys

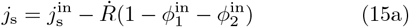

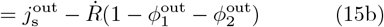

and thus we obtain the interface velocity expressed as the fluxes in the co-moving frame 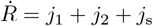.

We again relax the local equilibrium in the sharp interface model by considering different chemical potentials 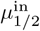 and 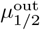 and osmotic pressures Π^in^ and Π^out^ at the interface. These differences give rise to the fluxes at the interface:

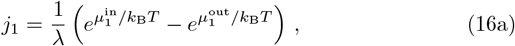

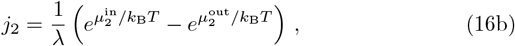

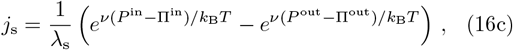

where we have the same kinetic coefficient *λ* for *j*_1_ and *j*_2_, since labeled and unlabeled molecules are considered to interact equally with the solvent at the interface.

We now calculate the boundary condition for the exchange of labeled and unlabeled material for a droplet that is otherwise at equilibrium, i.e.: 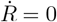 and *j*_1_ +*j*_2_ = 0, *j*_*s*_= 0 and *P* = Π. We define the equilibrium volume fraction profile as

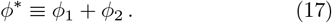

From Eq. (11a) and (11b) we obtain

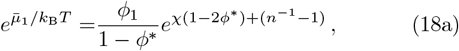

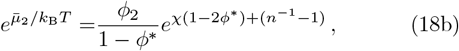

and thus

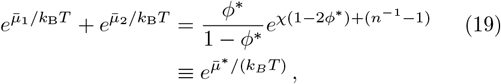

where we have defined

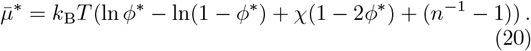

Since, at equilibrium *j*_1_ + *j*_2_ = 0, we have

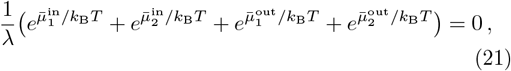

and 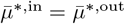. Using Eq. (19), therefore gives

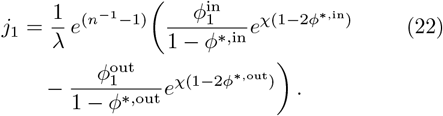

Defining the equilibrium partition coefficient

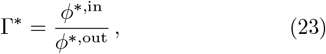

and the interface resistance

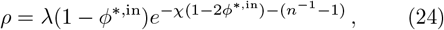

we obtain [15, 16]:

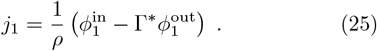

Eq. (25) describes both the thermodynamics of partitioning through a partition coefficient Γ^*∗*^ and interface resistance by the kinetic coefficient *ρ*.

### C. Relaxation to equilibrium

We numerically solve the dynamic Eq. (13a) for each phase with the boundary condition at the interface Eq. (25), and the flux relationships in the respective phases:

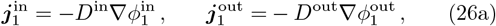

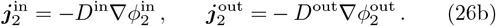

For a droplet at equilibrium, since 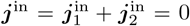, Eq. (26a) implies

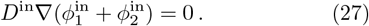

As initial condition, we use 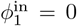 and 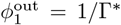. This initial condition is the situation after fully bleaching the droplets, i.e., the droplet contains only condensate molecules of type 2 (without a label). The following kinetics of exchanging the unlabeled molecules of type 2 with labeled molecules of type 1 in time corresponds to Fluoresecence Recovery after Photobleaching (FRAP) as observed experimentally (sketch in Fig. 2A and Refs. [3, 5, 6]). This recovery depends on the interface resistance *ρ* that is set by thermodynamic parameters and the kinetic coefficient *λ* (Eq. (24)).

The time course starts right after bleaching at time *t* = 0 (Fig. 2B). In the case for no interface resistance (*ρ* = 0), the equilibration occurs infinitely fast and the partition ratio reaches the partition coefficient immediately (dashed line). For non-zero interface resistance, the partition ratio approaches the partition coefficient asymptotically. This can also be seen in the time course of volume fraction profiles (Fig. 2C). As a result, gradients in the volume fraction of labeled molecules are weaker in the interface resistance case than the no-resistance case at comparable recovery stages. Since interface resistance is a kinetic phenomenon, the equilibrium volume fractions are identical.

Interestingly, interface resistance can also be modeled by using a finite-width interface-compartment [22], between the dense and dilute phases, which we present in Appendix A. This three-compartment model is of relevance, for example when interface resistance results from a wetted surface layer on condensate interfaces. In the Appendix A we show that the three-compartment model reduces to an effective interface resistance model in the limit of a narrow interface and when either of the following conditions is satisfied: (a) Small diffusion coefficient and finite partitioning into the interface. This corresponds to the mobility minimum scenario, which we discuss in Sect. III A. (b) Finite diffusion coefficient and small partitioning into the interface. This corresponds to the potential barrier case, discussed in Sect. III B.

## III. TRANSPORT ACROSS A CONTINUOUS INTERFACE

We now consider two continuous models that give rise to interface resistance. We focus on a case where the two phases are at equilibrium and transport is observed through labeling of molecules.

In such a binary mixture where some of the condensate molecules are labeled and others are unlabeled, the volume fraction of labeled molecules *ϕ*_1_ follows an effective diffusion equation [3, 4]:

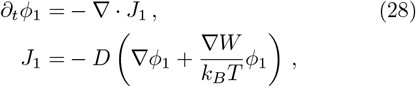

where *W* = − *k*_*B*_*T* ln *ϕ* is an effective potential and *D* = *k*_*B*_*T/γ* is the single-molecule diffusion coefficient, where *γ* is a friction that generally depends on volume fraction of condensate molecules *ϕ*.

For a system that phase-separates, the volume fraction changes from a high value inside the condensate *ϕ*^in^ to a low one outside the condensate *ϕ*^out^, for example in our reference case *ϕ*_0_ (grey line in Fig. 3E). A quantifier is the partition ratio

**Figure 3.**
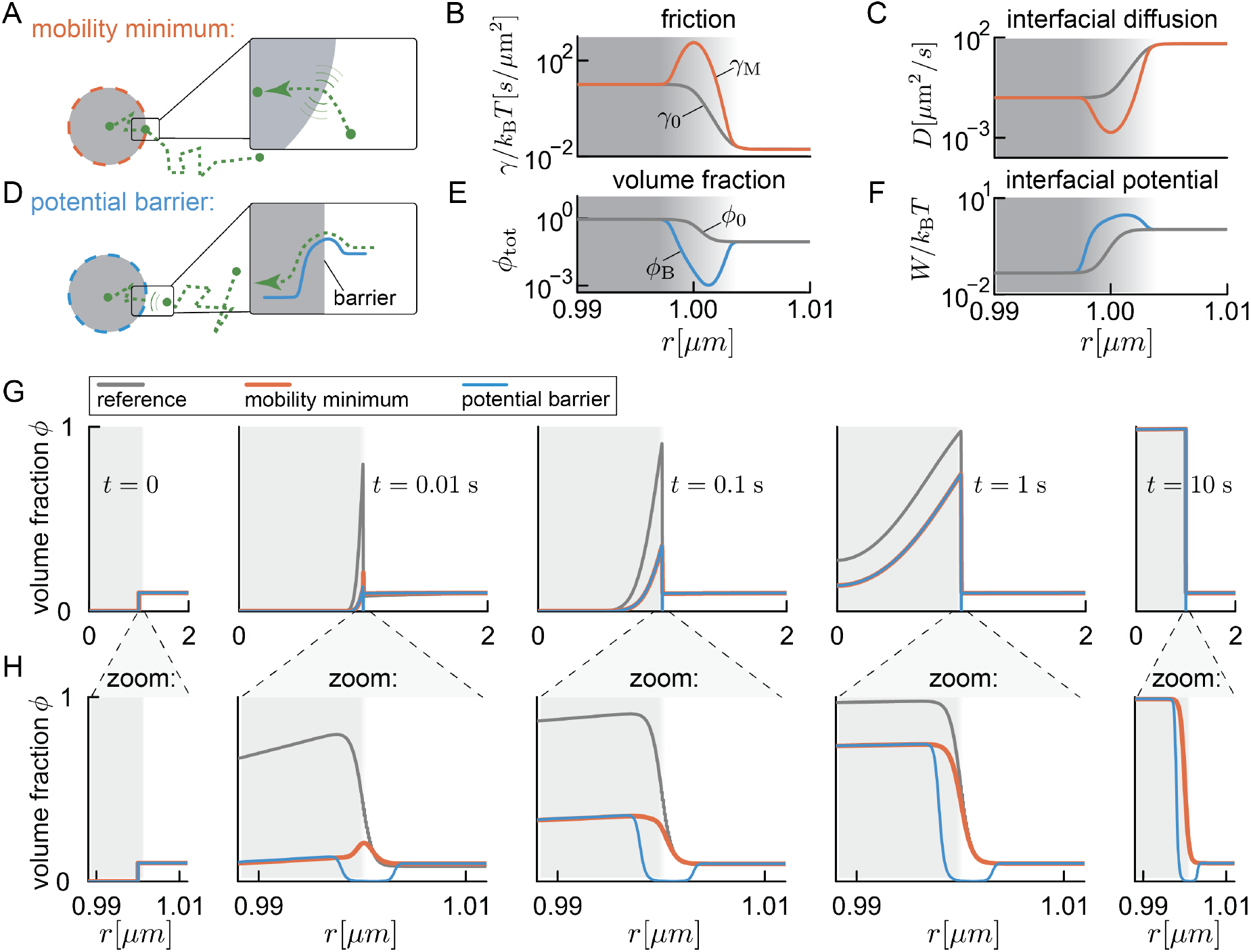
Continuous models of interface resistance. **(A)**: Interface resistance in continuous models can arise through a minimum in particle mobility at the phase boundary. **(B)**: If resistance is induced by a friction peak (orange, *γ*_M_), there is a pronounced mobility minimum and therefore in (C) a strongly reduced diffusion coefficient at the phase boundary. In the case of interface resistance via a potential barrier (grey), friction and diffusion coefficient interpolate monotonously between the values inside and outside of the phase boundary. **(C)** The friction peak at the phase boundary induces a corresponding minimum in the diffusion coefficient (orange). For the potential barrier case, the diffusion coefficient interpolates monotonously between dense and dilute phases (grey). **(D)** Interface resistance in continuous models can also arise through a potential barrier at the phase boundary. **(E)** In the case of interface resistance via a potential barrier, the volume fraction displays a pronounced minimum at the phase boundary (blue, *ϕ*_B_). **(F)** The minimum in (E) reflects a potential barrier at the phase boundary (blue). There is no potential barrier in the case of a mobility minimum (grey). **(G)** Time course of theoretical FRAP recovery for continuous boundary models. All models start with the same initial conditions. Away from the boundary, the concentration profiles of the mobility minimum and potential barrier cases are indistinguishable. **(H)** Zoom-in to the boundary region of panel (D). Note the distinct volume-fraction profiles within the boundary throughout recovery.

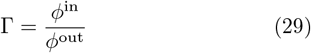

with its equilibrium value given by Eq. (23). If the interface is sharp, the change in condensate volume fraction *ϕ* occurs over a very short length scale. On large length scales, this effectively gives rise to a jump of the potential *W* across the interface. Using the formalism of Ref. [23], we can show how this potential jump creates a non-trivial boundary condition at the interface.

Let us consider the interface to be at *x* = *x*_0_. We call the position just left of the interface *x*^−^ and the one just right of the interface *x*^+^. Directly integrating Eq. 28 across the interface gives:

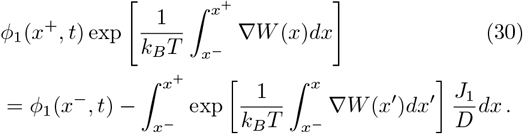

For a narrow interface, with finite *W*, *J*_1_, and *D*, the integral on the right-hand-side vanishes and we find

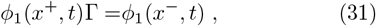

where we have used that the potential jump across the interface *W* (*x*^+^) − *W* (*x*^−^) = *k*_*B*_*T* ln Γ. Equation (31) is the quasi-equilibrium condition that was employed in Refs. [3, 4]. An interface resistance may arise if the integral on the r.h.s. of Eq. (30) does not vanish. Recalling that *W* = −*k*_*B*_*T* ln *ϕ*, from Eq. (30), we obtain

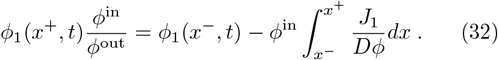

For a finite flux *J*_1_, the integral on the r.h.s. does not vanish for a small diffusion coefficient *D* or a small value of *ϕ* at the interface. We consider two ways to realize each of these cases: (A) a lowered diffusivity and (B) a potential barrier at the interface.

### A. Mobility minimum at the Phase Boundary

To obtain a lowered diffusivity at the interface, we introduce an increased friction

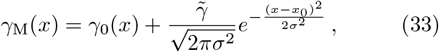

at the position *x*_0_ of the interface, where *γ*_0_ is the reference friction (grey line in Fig. 3B) and *σ* is the width of the high-friction region (Fig. 3A,B). The peak in friction corresponds to a reduced diffusion coefficient *D* at the phase boundary since *D* = *k*_*B*_*T/γ*_M_ (Fig. 3C). This translates to a slower transport of labeled molecules across the phase boundary compared to the reference case without enhanced interface resistance (orange traces in Fig. 3G). Note the transient accumulation of labeled molecules at the phase boundary at early times (Fig. 3H, local maximum of orange curve at *t* = 0.01 s), which vanishes at equilibrium when the profiles of the reference case and the case with a mobility minimum are identical.

The case of a mobility minimum at the interface can be discussed in the sharp interface limit, by taking the limit *σ →* 0 in Eq. (33)^1^. The friction profile in Eq. (33) then becomes approximately

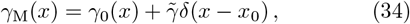

which makes the boundary condition at the interface (Eq. (32))

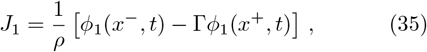

where

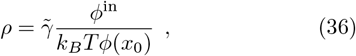

is the interface resistance. Eq. 35 is the effective boundary condition used in Ref. [5] and equivalent to Eq. 25.

### B. Potential Barrier at the Phase Boundary

Next, we consider interface resistance caused by a potential barrier at the interface (Fig. 3E, F). To achieve the same interface resistance as in the mobility minimum case (Eq. (36)), we need the denominator of the integral in Eq. (32) to be equal between both cases. We achieve this by setting

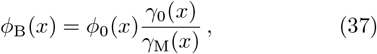

where *ϕ*_0_(*x*) is the reference profile without the potential barrier, *γ*_0_(*x*) is the reference friction profile without resistance, and *γ*_M_(*x*) is the friction profile with a peak given in Eq. (33). The friction coefficient for this case, follows the profile without the peak, *γ*_0_(*x*) (grey line in Fig. 3B). The effective potential corresponding to Eq. (37), *W*_*B*_ = − *k*_*B*_*T* ln *ϕ*_*B*_, is peaked at the phase boundary (Fig. 3F):

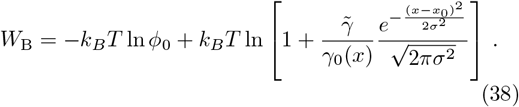

The resulting peak in the potential corresponds to a minimum of the volume fraction (Fig. 3E). The dynamics far from the interface can thus be matched with the case of a mobility minimum (overlap of blue and orange in Fig. 3G,H, for a brief discussion of the numerics see appendix D). Notably, the minimum volume fraction is also visible at equilibrium, distinguishing it from the mobility minimum case (Fig. 3H).

## IV. QUANTIFICATION OF INTERFACE RESISTANCE

Current techniques to measure interface resistance rely on quantifying the exchange of fluorescently labeled molecules across a stationary phase boundary (see Refs. [5, 6, 12], and Fig. 2 for an example of FRAP). Quantifying the FRAP halftime for different interface resistance *ρ* shows that in particular for small droplets, the recovery time is dominated by interface resistance (large spread of halftime for small radii in Fig. 4A). For larger droplets the diffusive time scales become dominant, as discussed in depth in Ref. [6].

**Figure 4.**
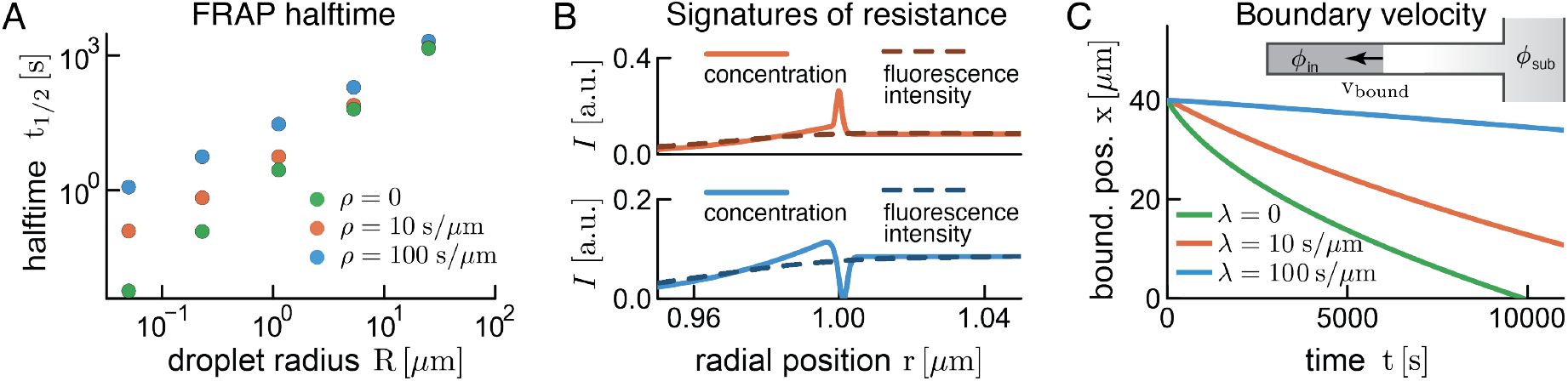
Quantification of interface resistance with different experimental techniques. **(A)**: FRAP halftime depends sensitively on interface resistance, particularly for small droplets. For larger droplets, diffusive time scales become dominant (see discussion in [6]). **(B)**: Given the current estimates of interfacial width, direct observation of phase boundary concentrations (orange and blue curves) via ensemble fluorescence microscopy (intensity *I*) is not possible due to the much larger PSF. **(C)** Suggested microfluidic assay to measure interface resistance. A capillary containing the dense and dilute phases is connected on the dilute side with a sub-saturated channel. All numerical solutions have *λ* = *λ*_s_.

Alternatively, one could try to directly observe the signatures of interface resistance at or within the boundary, similar to the theoretical magnification of boundary dynamics in Fig. 3H. Early time points provide the best opportunity for such measurements, as the concentration peak (mobility minimum case) or the concentration minimum (potential barrier case) are most prominent compared to the surrounding fluorescence of the adjacent phases. However, even during early times such measurements are impossible with current techniques for samples with a large fraction of labeled molecules (Fig. 4B), due to the large point-spread function (*>* 250 *μm*) compared to the interface width seen in experiments and simulations (*<* 10 *μm*) [24, 25]. Assuming such a 25fold difference we see a completely smoothed-out interface (dashed lines in Fig. 4B), where the dashed lines become hardly distinguishable. However, recent progress in single-molecule imaging [26] enables nanometer resolution imaging, and thus provides an exciting opportunity to study phase boundaries in detail.

As labels are not always available or desirable [18] an alternative for measuring interface resistance could be provided by microfluidics as depicted in Fig. 4C. Briefly, a channel that contains the dense phase is connected on one side to a fluid channel below the saturation concentration (*ϕ*_sub_ in Fig. 4C), which effectively keeps the boundary concentration constant. Due to diffusive material exchange, the dense phase will shrink, and the resulting speed of the phase boundary will depend on the interface resistance. Here, the most robust read-out can be obtained at long times, provided the experimental assay is sufficiently stable. We note that the dynamics are sensitive to the experimental set-up, in particular with respect to channel length and outside concentration *ϕ*_sub_. Generally, a large concentration difference seems desirable between the outside reservoir and the capillary containing the dense phase.

## V. CONCLUSIONS

In this work, we introduce a theoretical framework to study the kinetics of molecules across an interface between two coexisting liquid phases. Molecule transport can lead to movement of this interface and to material exchange between the two phases. We started by deriving the interfacial kinetics in the thin-interface limit. Crucially, we allow for a discontinuity in the chemical potential at the interface, which naturally leads to interface resistance, as proposed for droplet growth [16] and particle exchange across liquid-liquid interfaces [5, 6, 12]. Our theory extends to multiple molecular species. We show that interface resistance leads to a slow-down in interfacial movement in a binary mixture, and, if part of the molecules are labeled, a decelerated molecule exchange between labeled and unlabeled molecules across the phase boundary. We then present continuous transport models that introduce interface resistance, either by slowing down diffusion at the interface, or by introducing a potential barrier at the interface.

Our theory comes at a time when transport across phase boundaries is receiving renewed interest, both in biology and engineering. For example, a number of works have described how client molecules that are not involved in phase separation travel across a stationary liquidliquid phase boundary in aqueous two-phase systems in an external electrical field [7, 8, 12, 27, 28]. A transient accumulation of material at the interface was observed. This accumulation is reminiscent of interface resistance, and has been attributed to the Donnan potential across the interface and its interaction with the electric field that was applied across the interface. This interplay was suggested to cause a local potential well, where molecules are trapped. This corresponds to the opposite of the potential barrier case discussed here. A main difference is that in the experiments, due to the electric field, no equilibrium distribution comparable to ours can be observed – particles eventually desorb from the boundary and travel across the entire system.

For biomolecular condensates, it was suggested that interface resistance is required to explain FRAP recovery curves of the protein LAF-1 [5]. Taylor et al. found that the slow recovery of condensates could not be accounted for based on the measured diffusion and partition coefficients. By introducing an interface resistance parameter the discrepancy was resolved. However, adding this new parameter to the model led to a reduced fit quality (see Fig. 11 in Ref. [5]), indicating that these fits do not provide sufficient evidence to conclude that interface resistance governs the FRAP recovery of LAF-1. Using the available spatio-temporal resolution of the data could improve the parameter quantification [3].

We suggest a number of ways to measure interface resistance despite current optical limitations in resolving the phase boundary. Similar to the FRAP measurements performed in Ref. [5], we suggest measuring the FRAP halftime. However, ideally this would be done for differently sized condensates and by also using the spatial information available, instead of integrating the intensity over the entire droplet. This has the advantage of directly measuring *D*_in_, which avoids any confusion between the time scale associated with the interface and that associated with diffusion inside the droplet. Even without labeled molecules, interface resistance could be measured by examining droplet growth, for example using a microfluidic setup as proposed in Fig. 4C. In conclusion, our framework can guide further experiments to quantify interface resistance and decipher the underlying molecular mechanisms.

## ACKNOWLEDGEMENTS

We would like to thank Patrick McCall, Giancarlo Franzese and Arghya Majee for insightful discussions.

## Appendix A Interface compartment model

The interface compartment extends from position − *a* to *a* and we denote the space-dependent interface volume fraction profile as 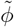, and the constant diffusion coefficient in the interface compartment as 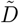. We have a diffusion equation in each compartment

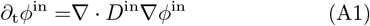

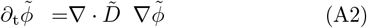

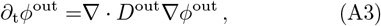

which are coupled by non-trivial boundary conditions where the compartments meet (at −*a* and *a*)

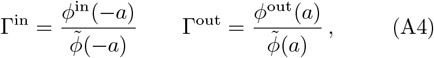

where we have introduced the two partition coefficients, which describe how enriched the dense and the dilute phases are with respect to the interface at the boundary, respectively. For a narrow interface *a ≪* 1 we assume that the interface compartment rapidly reaches a steady state, characterized by a constant flux in space 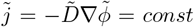. This constant flux implies a linear volume fraction profile inside the interface compartment. Continuity then implies that, at the boundaries between the interface compartment and the two phases, we have 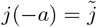 and 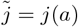, so that

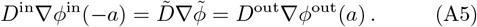

The linearity of the volume fraction profile in the interface compartment allows us to write

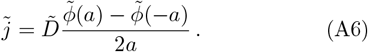

Combining this expression with Eqs. (A4) and (A5), we have that

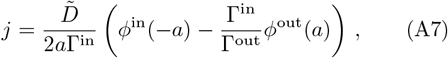

which, identifying,

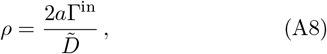

corresponds to the interface resistance boundary condition obtained for the effective droplet model in Eq. (25) if we define 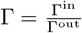. Since *a* is small we need to require the ratio 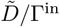 to be small as well to obtain a finite interface resistance *ρ*. This can be achieved by having a small diffusion coefficient 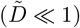, which is what we discussed in Section III A for the case of a mobility minimum at the interface. A finite interface resistance can also arise for large partition coefficients enriching the two phases with respect to the interface compartment (Γ^in^ *≫* 1 and Γ^out^ *≫* 1, with finite Γ). This latter case corresponds to a potential barrier in the interface, as discussed in Section III B.

## Appendix B Numerical solution of a moving boundary in a binary mixture

The solutions shown in Fig. 1B-D and Fig. 4C are found using a custom finite difference scheme implemented in Matlab. Code is available in the corresponding Repository.

## Appendix C Numerical solution of exchange of labeled and unlabeled molecules

The solutions shown in Fig. 2 and Fig. 4A were obtained using a customized Crank-Nicolson scheme implemented in Julia, available in the corresponding repository.

## Appendix D Numerical solution of the continuous interface models

To solve Eq. 28, we use a custom Matlab implementation available at the corresponding repository. Briefly, for the solutions in Fig. 3G,H for the mobility minimum case we use a friction of the form

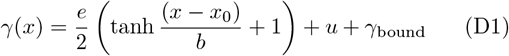

where we use *γ*_bound_ = *f* exp(− (*x* − *x*_0_)^2^*/b*^2^) in the case of the mobility minimum and *γ*_bound_ = 0 in the case of a potential barrier.

The volume fraction is of the form

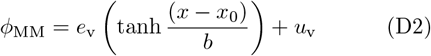

in the case of the mobility minimum, and of the form Eq. 37 in the case of the potential barrier. Parameter values can be obtained from the corresponding repository.

## Footnotes

1 The width of the low mobility region, *σ*, needs to be smaller than the interface region. If *σ* is as large as the interface, one still obtains interface resistance as described in Eq. (35), but the value of the resistance *ρ* will be different and depend on the shape of the volume fraction profile inside the interface region.

